# Hormone gene expression changes in the zebrafish caudal neurosecretory system under acute environmental challenges

**DOI:** 10.1101/2025.01.12.632590

**Authors:** Bérénice Bichon, Gladys Alfama, Anne-Laure Gaillard, Hervé Tostivint, Guillaume Pézeron

## Abstract

The caudal neurosecretory system (CNSS) is a neuroendocrine complex, found only in fishes, suggested to be involved in osmoregulation and thermo-adaptation. Although being discovered 70 years ago, the CNSS has been understudied and its physiological significance remains poorly understood. The zebrafish (*Danio rerio*) is a well-established model organism for functional studies, yet, so far, the role of the CNSS has not really been investigated in this species. In the present study, we attempted to identify environmental factors whose variations induce changes in CNSS activity. Juvenile zebrafish were submitted to acute (2 h, 8 h, and 24 h) pH, salinity, and temperature challenges, and CNSS activity was quantified by measuring the expression levels of peptide hormone-encoding genes using quantitative PCR. Our findings revealed that temperature challenge affected the expression of the largest number of gene, followed by salinity and pH challenges. This suggested that, in the zebrafish, the CNSS is involved in thermo-adaptation and osmoregulation, which is consistent with what has been reported in other species, and showed that the zebrafish is a relevant model to study the function of the CNSS.

## 1 Introduction

Facing numerous environmental variations, potentially inducing stress, fish are compelled to implement adaptive strategies to preserve homeostasis. In animals, maintaining physiological balance relies on mechanisms that buffer out external variations to maintain stable internal conditions. In vertebrates, one of the key players in maintaining homeostasis is the hypothalamo-hypophyseal neuroendocrine system, which plays a central role for adapting to environmental variations by orchestrating appropriate physiological responses.

Fishes possess a second neuroendocrine complex located in the caudal part of the spinal cord, known as the caudal neurosecretory system (CNSS) (Rousseau et al., 2024). First described in the Japanese eel (*Anguilla japonica*) (Enami, 1955), the CNSS exists in all groups of jawed vertebrates (Gnathostomata) studied to date, except sarcopterygians (Enami and Imai, 1955). In teleosts, the CNSS consists of a small population of large secretory neurons, known as Dahlgren cells (Dahlgren, 1914; Speidel, 1922), that project their non-myelinated axonal tracts toward a highly vascularized ventral expansion of the spinal cord called urophysis (Arnold-Reed et al., 1991; Bern, 1985; Fridberg and Bern, 1968). In this neurohaemal organ, secretory products are stored within axonal swellings of Dahlgren cells before being released into the bloodstream. Urophysial capillaries merge to form the caudal vein, which ensures its drainage and joins the portal-renal system. Beyond the kidney, the secretion products of the CNSS then access the systemic circulation, potentially interacting with renal cells, interrenal tissue, and the Corpuscles of Stannius (Bern et al., 1985).

The CNSS produces several peptide hormones, synthetized by Dahlgren cells and released by the urophysis. The two main secretion products of the CNSS are Urotensin 1 (Uts1; also known as UI, and Urocortin 1 in mammals) and Urotensin 2 (Uts2; also known as UII). Both were characterized from urophysial extracts on the basis of their pharmacological activities. Uts1 is related to the Corticotropin-releasing hormone (Cardoso et al., 2016; Lovejoy and de Lannoy, 2013) and exhibits blood pressure-lowering properties (Lederis et al., 1982). Uts2 is a cyclic neuropeptide, phylogenetically related to Somatostatin (Tostivint et al., 2013), that induces blood pressure elevation and smooth muscle contraction (Pearson et al., 1980). Additional peptide hormones were shown to be produced by the CNSS in different species. These include Corticotropin-releasing hormone (Crh) (Lu et al., 2004), Parathyroid hormone like hormone (Pthlh, also known as Parathyroid hormone-related protein) (Ingleton et al., 2002; Lu et al., 2017), Met-enkephalin (Yamada et al., 1988), and Staniocalcin (Greenwood et al., 2009). Further, Arginine-vasopressin (Avp, also known as Vasotocin) and Oxytocin (Oxt, also known as Isotocin) were found in the urophysis in different teleosts, but whether these hormones are actually produced by the Dahlgren cells remains uncertain (Gozdowska et al., 2013; Lacanilao, 1972).

Since its first description, the CNSS has been proposed to play a role in fish adaptation to environmental changes, in particular in osmoregulation (Enami et al., 1956). Consistent with this idea, Uts1 and Uts2 were reported to modify ionic fluxes through skin (Marshall and Bern, 1981, 1979), intestine (Loretz et al., 1983; Mainoya and Bern, 1982) and urinary bladder (Loretz et al., 1983). Moreover, CNSS activity is affected upon salinity challenge in different species. In the rainbow trout (*Oncorhynchus mykiss*), transfers from freshwater (FW) to seawater (SW) resulted in an increase of Uts1 immunoreactivity in Dahlgren cells but a reduction in the urophysis, suggesting higher secretion in the bloodstream (Larson and Madani, 1996). Similarly, increased levels of *uts1* and *crh* mRNA in caudal spinal cord were also reported following transfer to SW (Craig et al., 2005). As for Uts2, following transfer to SW, its secretion was transiently reduced (2 h after transfer), but was increased upon longer exposition (Larson and Madani, 1996). In the European flounder (*Platichthys flesus*), transfer from SW to FW increased *uts2* mRNA levels in the CNSS, while lowering plasma Uts2 concentration (Bond et al., 2002; Lu et al., 2006). In the same condition, *uts1* expression was transiently increased (8 h after transfer), but the urophysial content of Uts1 was reduced (Lu et al., 2019).

More recently, the CNSS was also suggested to be involved in thermo-adaptation. In the olive flounder (*Paralichthys olivaceus*), hypothermia led to increased *crh* and *uts2* mRNA levels in the CNSS, while hyperthermia increased numbers of Uts1-positive Dahlgren cells and *uts1* mRNA levels (Yuan et al., 2019). Further, in this species, the CNSS was found to directly respond to hypothermal challenge through Transient Receptor Potential (TRP) channels (Yuan et al. 2020), and this response seems to differ according to the temperature and the duration of the challenge (Yuan et al., 2021). Yet, in the rainbow trout, by contrast, cold stress did not induce any change in CNSS mRNA levels of *crh* and *uts1* (Craig et al., 2005).

The role of the CNSS was also studied through urophysis ablation, but this led to inconsistent results. In some cases, alterations in ion balance were reported (Chester Jones et al., 1969; Turtle, 1974), while in others, no effects were observed (Baldisserotto et al., 1994; Fryer et al., 1978; Imai et al., 1965). Therefore, the physiological significance of the CNSS remains poorly understood. The zebrafish is a well-known teleost model with unique advantages for functional studies. However, to date, only two studies focused on the function of the CNSS in this species. One study reported that exposure of zebrafish embryos and young larvae (from 2 to 6 days post-fertilization, dpf) to SW increased *uts2a* and *uts2b* expression at 6 dpf (Luo et al., 2014). Another study found that exposure to cold or hot stress downregulated the expression levels of *crh, uts1, uts2a* and *uts2b* (Luo et al., 2014). However, as these experiments were conducted at very early developmental stages, at which the urophysis is probably not yet formed (Fridberg and Bern, 1968), the significance of these variations remains difficult to evaluate. In the present study, in order to determine the physiological responses in which the CNSS plays a role in the zebrafish, we have attempted to identify environmental variations that induce changes in CNSS activity. We have investigated the effects of acute changes in water pH, salinity and temperature on the expression levels of peptide hormone-encoding genes in the CNSS. Our findings suggested that, in the zebrafish, the CNSS is involved in osmoregulation and thermo-adaptation, and showed that the zebrafish is a relevant model to study the functions of the CNSS.

## 2 Materials and methods

### 2.1 Animals

Zebrafish (*Danio rerio*) used in this work were wildtype (Tübingen strain, from EZRC https://www.ezrc.kit.edu). All animals were bred in our facility and maintained according to standard procedure (Westerfield, 2007). Zebrafish embryos were obtained by natural spawning, collected from breeding tanks and kept in E3 embryo medium until 5 dpf. Larvae between 5 and 15 dpf were raised in water containing 5 g/l of sea salt (Instant Ocean) and fed *ad libitum* with rotifers. Both embryos and larvae were maintained in an incubator at 28.5°C under a 14-hour light/10-hour dark cycle. From 15 dpf onward, fish were kept in an Aquaneering standalone system with water at 26-27°C, pH 7-8, and conductivity 200-300 µS. Animals were maintained under a 14-hour light/10-hour dark cycle and fed once a day with live artemia and twice a day with dry food (Gemma Micro ZF, Skretting).

### 2.2 Ethic approval

This study was conducted in accordance with the European Communities Council Directive (2010/63/EU) and all procedures were approved by the Cuvier Institutional Ethics Committee at the Muséum national d’Histoire naturelle (APAFIS#35948). The study aimed to follow ARRIVE guidelines (Sert et al., 2020).

### 2.3 Experimental design for environmental challenge

The procedure was designed to assess the impact of acute environmental variations. Three water parameters were tested: temperature, pH and salinity. For each parameter, three conditions (low, control and, high) and three time points (2 h, 8 h, and 24 h) were used.

Assays were conducted as follows: on the evening prior to the experiment, 63 sibling juvenile zebrafish (10 weeks, 15-18mm standard lenght(Parichy et al., 2009)) were randomly distributed into 9 groups, and placed in 1000 ml beakers (7 fish per beaker) containing 500 ml of control water (see next section). Fish were kept overnight in an incubator at 26°C with the same light-dark cycle as the facility. On the following morning, within one hour after lights turn on, the control water was replaced with high-condition water (see next section) for three beakers, low-condition water (see next section) for three other beakers, and control water for the last three. Then, fish from one beaker for each water condition (high, low and control) were collected 2 h, 8 h, and 24 h after the initial water change. At each time point, fish were rapidly euthanized using anesthetic overdose (tricaine methanesulfonate - MS222, 0.03% in high, low, or control water) and sample were collected (see below). To maintain optimal water quality for the 24 h time point, the water was replaced with new high, low or control water before the second night.

### 2.4 Preparation of conditioned waters

In each experiment, the control water was prepared with deionized water supplemented with sea salt (Instant Ocean) at 150 ppm (0.15 g/l, conductivity ∼250µS). The pH was maintained to 7.5 with HEPES (1 M) and the temperature was kept at 26°C.

For pH challenge, the high-condition water was at pH 9.0 and the low-condition water was et pH 6.0. The conditioned waters were obtained from control water with the pH adjusted with HCl or NaOH. For the salinity challenge, the high-condition water was 5000 ppm (5 g/l) and the low-condition water was 5 ppm (0.005 g/l, ∼15 µS). The conditioned waters were obtained from control water by adding sea salt or by dilutiing with deionized water. Finally, for temperature challenge, the high-condition corresponded 34°C and the low-condition 18°C. Here, control water was simply placed into different incubators set to the corresponding temperatures.

### 2.5 Sample collection

Following euthanasia, fish were dissected in ice-cold PBS. The anterior part of the fish was removed by cutting at the posterior end of the anal fin, and the caudal fin was removed by cutting at its base. Samples were placed individually in vials and snap-frozen in liquid nitrogen. All samples were stored at -80° C before further processing.

### 2.6 RNA extraction and real time quantitative RT-PCR

Total RNAs from samples were extracted using TRIzol Reagent (Invitrogen). Samples were directly immersed in 500 μL of TRIzol reagent, and homogenized using a TissueLyser II system (Qiagen). Following RNA extraction, samples were treated with DNase I (Roche) to remove potential contaminating genomic DNA, and then purified using phenol/chloroform extraction. The integrity of RNA was verified by electrophoresis on 1% agarose gel, and RNA concentration was determined using a NanoDrop 2000 spectrophotometer (Thermo Fisher). Then, cDNAs were obtained using 500 ng of RNA with GoScript reverse transcriptase (Promega) and random primers (Promega). cDNAs were diluted 10-fold prior to real-time PCR. Quantification was carried out by real-time PCR using specific primer pairs and PowerUp SYBR Green (Applied Biosystems) on a QuantStudio™ 6 Flex Real-Time PCR System (Applied Biosystems). Primer pairs were validated on 4 serial dilutions of corresponding cDNA. Linearity of CT variation on dilutions and single peak of melting curved were verified. All primer sequences are indicated in Table S1. For all quantitative RT-PCR (RT-qPCR) experiments, reference genes were *lsm12b*, *mob4* (Hu et al., 2016) and *tdr7* (Vanhauwaert et al., 2014). Fold changes were calculated using the 2e-ΔΔCT formula.

### 2.7 Statistics and figure preparation

Prior to statistical analysis, outliers in gene expression data from RT-qPCR were identified and removed using interquartile range (IQR) method. Specifically, for each gene, and within each time and each condition, samples with value more than 1.5 IQR below first quartile (Q1) or more than 1.5 IQR above third quartile (Q3) were considered outliers and excluded from data. Then, for each gene, and within each time and each condition, statistical analysis was performed using two-sample Wilcoxon tests to compare values from high and low condition to the control.

Analyses were performed using the R software (R Core Team, 2021) along with the Tidyverse package (Wickham et al., 2019). Graphs were created using the ggplot2 package (Wickham, 2016). Figures were prepared using Scribus software (www.scribus.net/) and Microsoft PowerPoint (Microsoft Corporation).

## 3 Results

In fish, the caudal neurosecretory system (CNSS) is thought to be involved in adaptation to environmental changes (Rousseau et al., 2024). In this study, we investigated whether acute changes in water pH, salinity, or temperature resulted in modification of CNSS gene expression levels in zebrafish. For this, juvenile fish were exposed to three experimental conditions (control, high-condition, and low-condition) for each parameter and samples were collected after 2 h, 8 h, and 24 h (Fig.1). In all cases, samples were used for RNA extraction and gene expression levels were measured using quantitative PCR. Genes assayed in this work (listed in Table 1) encode neuropeptides previously reported to be produced by the CNSS (Rousseau et al., 2024). Note that we could not obtain specific primers for *stc1* nor *stc1l* and thus these two genes were not included in this work. Also, in all assays, the expression levels for *avp* and *crha* were too low to be analyzed and the corresponding data are not presented.

**Fig 1.**
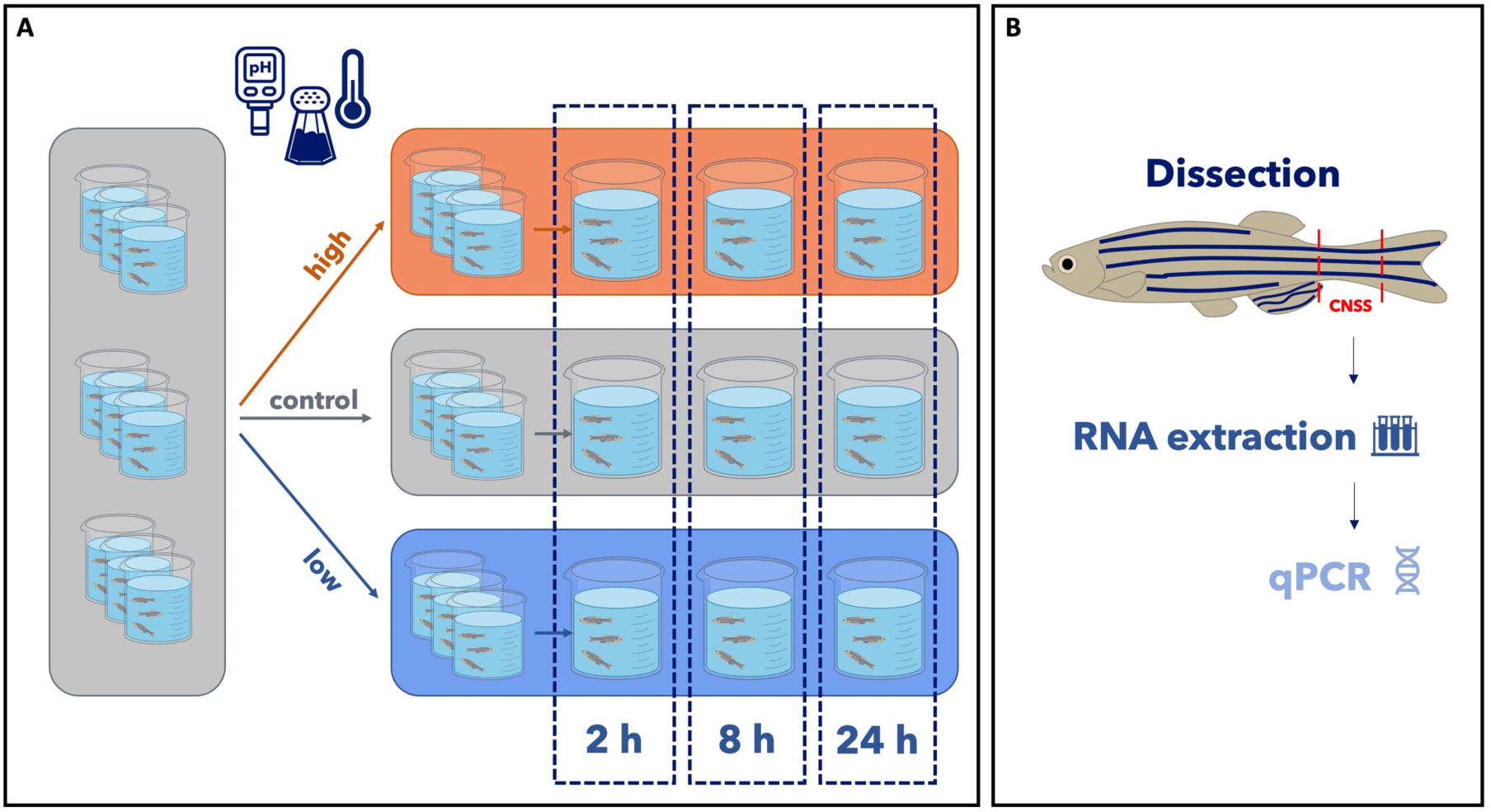
Overview of the experimental design. (A) The day prior to the experiment, juvenile zebrafish were randomly distributed into nine beakers with control water (7 fish per beaker). On the next morning, water was replaced with high, low or control condition water (see also method section). Samples were collected after 2 h, 8 h and 24 h. (B) Fish were dissected and the caudal parts (minus the tailfin) were retrieved and processed for quantitative PCR analysis.

**Table 1.** List of genes analyzed in this study. The ZFIN ID indicates the gene ID in the Zebrafish Information Network (ZFIN) database.

| Gene name | Symbol | ZFIN ID |
| --- | --- | --- |
| *arginine vasopressin* | *avp* | ZDB-GENE-030407-2 |
| *corticotropin releasing hormone a* | *crha* | ZDB-GENE-070912-371 |
| *corticotropin releasing hormone b* | *crhb* | ZDB-GENE-041114-75 |
| *oxytocin* | *oxt* | ZDB-GENE-030407-1 |
| *proenkephalin a* | *ppenka* | ZDB-GENE-030729-31 |
| *proenkephalin b* | *ppenkb* | ZDB-GENE-030729-3 |
| *parathyroid hormone-like hormone a* | *pthlha* | ZDB-GENE-050119-9 |
| *parathyroid hormone-like hormone b* | *pthlhb* | ZDB-GENE-110411-206 |
| *stanniocalcin 1* | *stc1* | ZDB-GENE-060825-79 |
| *stanniocalcin 1, like* | *stc1l* | ZDB-GENE-040426-1521 |
| *stanniocalcin 2a* | *stc2a* | ZDB-GENE-041008-14 |
| *stanniocalcin 2b* | *stc2b* | ZDB-GENE-120203-5 |
| *urotensin 1* | *uts1* | ZDB-GENE-041014-348 |
| *urotensin 2, alpha* | *uts2a* | ZDB-GENE-040630-11 |
| *urotensin 2, beta* | *uts2b* | ZDB-GENE-040630-10 |

### 3.1 Effects of pH challenge

Fish were exposed to water at pH 6 (low condition), pH 7.5 (control condition), and pH 9 (high condition). Figure 2 depicts the impact of these conditions on gene expression levels. In the 2 h challenge, *penka* was the only gene with modified expression levels, showing a clear reduction in the high condition compared to the control condition. At 8 h, the same tendency was observed for *penka* and two other genes had modified expression levels: *crhb* and *stc2b* with a reduction in the low condition. At 24 h, three genes had modified expression levels: *oxt* was increased in low condition, *stc2b was* reduced the high condition and *uts2b* was reduced in both low and high conditions. Thus, overall, the pH challenge resulted in limited changes in gene expression.

**Fig 2.**
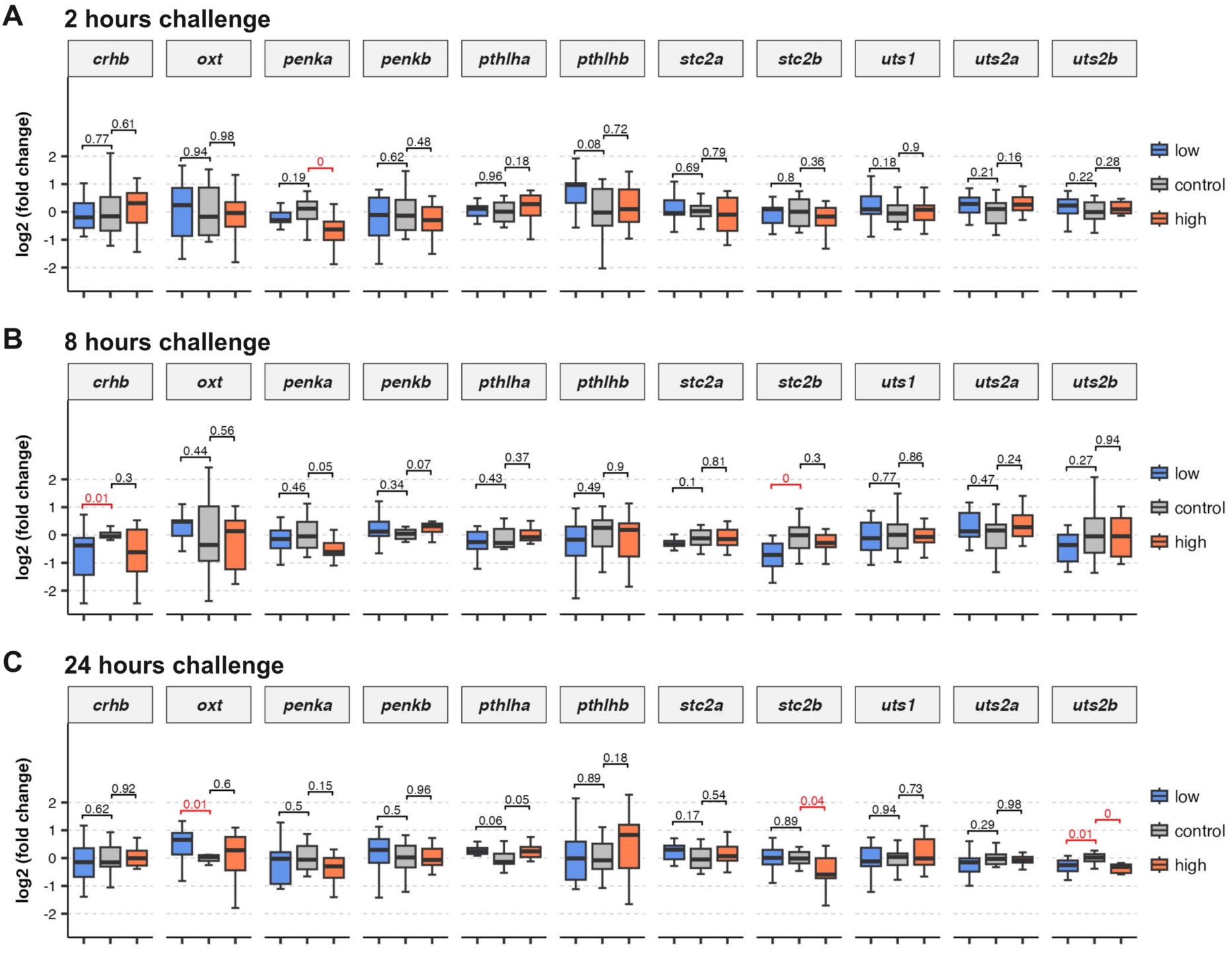
Gene expression changes following acute pH challenge. Expression levels of indicated genes after 2 hours (A), 8 hours (B) and 24 hours (C) pH challenge with low, control and high conditions. In all cases, results are expressed as log2 of fold change (ΔΔCT) compared to the control condition.

### 3.2 Effects of salinity challenge

Fish were exposed to water with salts at 5 ppm (low condition, see also methods), 150 ppm (control condition), and 5000 ppm (high condition). Gene expression levels under these conditions are illustrated in Figure 3. After 2 h, two genes, *oxt* and *uts1*, showed significant changes in expression levels compared to the control. o*xt* expression was increased in the low salinity condition, with a similar tendency in high condition (p = 0.06), and *uts1* exhibited higher expression level in the high condition. With the 8 h challenge, we found that the expression level of *penka* was increased in the low condition. Consistent with our observation in the 2 h assay, our result suggested an increase of o*xt* expression level in the high condition (p = 0.08). Additionally, we also observed an increase in *uts2b* expression levels under the high condition (p = 0.05). Finally, in the 24 h challenge, we observed an upregulation of *stc2b* expression levels in the low salinity condition. We also noted a marked increase (approximately 2-fold) in *uts2b* expression levels in the low condition compared to the control. Interestingly, with the high condition, our results showed increased expression levels for *uts2a* and suggested upregulation of *uts2b* (p = 0.05), and *uts1* (p = 0.07) as well. In conclusion, challenging zebrafish with salinity variations had limited effect on gene expression. Yet, it appeared to affect the expression levels of *uts1, uts2a* and *uts2b* suggesting that, in zebrafish, Uts1 and Uts2 could be involved in the response to salinity variation as previously suggested by studies in various teleost species (Rousseau et al., 2024).

**Fig 3.**
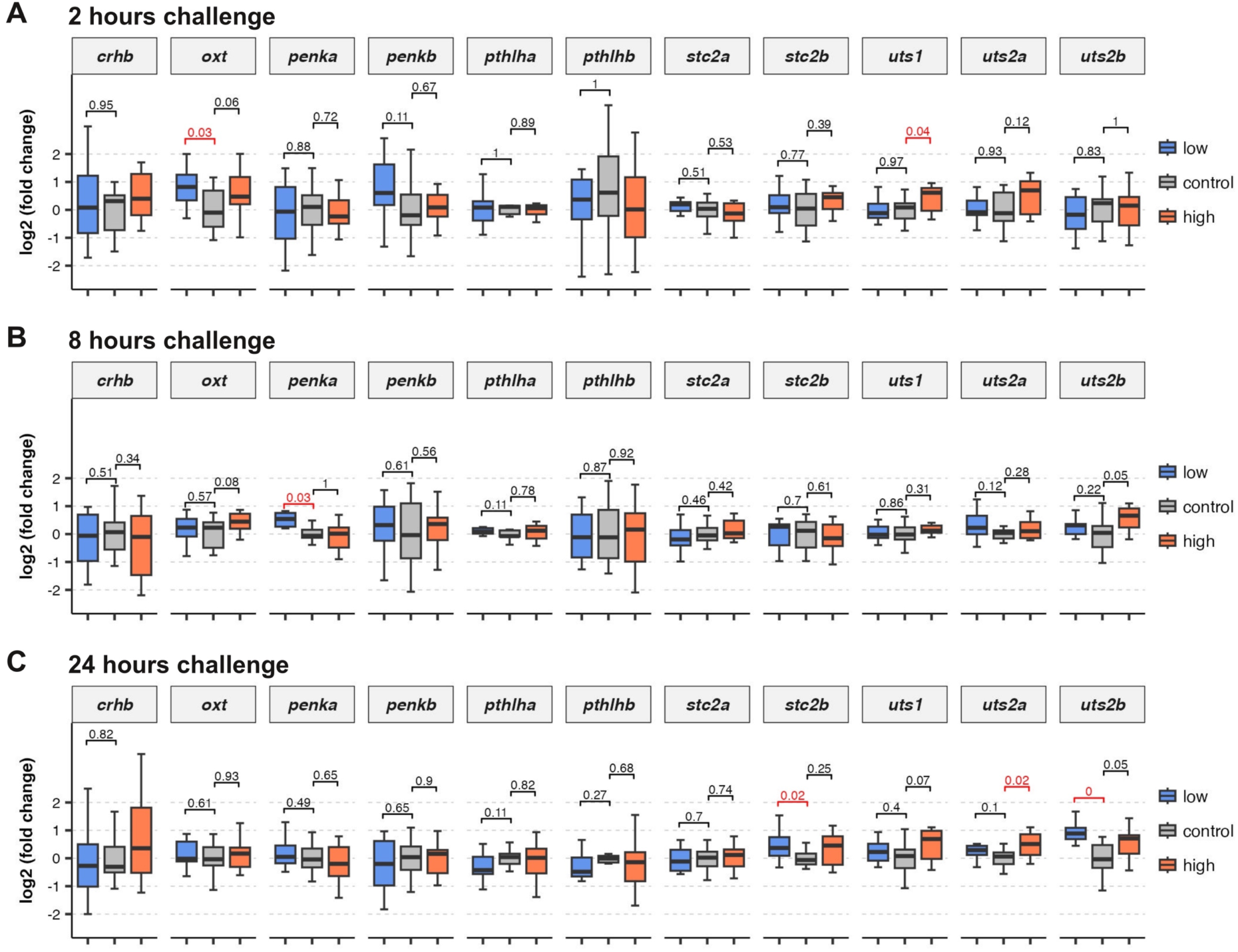
Gene expression changes following acute salinity challenge. Expression levels of indicated genes after 2 hours (A), 8 hours (B) and 24 hours (C) salinity challenge with low, control and high conditions. In all cases, results are expressed as log2 of fold change (ΔΔCT) compared to the control condition.

### 3.3 Effects of temperature challenge

Fish were exposed to water at 18° C (low condition), 26° C (control condition), and 34° C (high condition). The effects on gene expression levels are presented Figure 4. In the 2 h challenge, the expression levels of *penka* and *stc2a* were reduced in the high condition. This was also the case for *pthlha* in the low condition and, to a lesser extent, in the high condition (p = 0.05). Conversely, our data suggested increased expression levels of *stc2b* in the low condition (p = 0.05). Also, both *uts1* and *uts2a* expression levels appeared clearly increased in the high condition compared to the control. At 8 h, most gene expression levels were altered in either the low or the high condition. In the low condition, *pthlha*, *pthlhb*, and *stc2a* were down-regulated compared to the control. This seemed to also be the case for *oxt* (p = 0.05). In the high condition, we observed increased expression levels for *crhb* and *uts2a*, and reduced expression levels for *oxt, penka*, *penkb* (p = 0.07). For the 24 h challenge, *penka*, *penkb, stc2a* and *stc2b* appeared downregulated in the high condition, whereas, in the low condition, we observed an upregulation for *oxt*, *pthlha*, *stc2a* and *crhb* (p = 0.05). Finally, the expression of *uts1* was increased in the low temperature condition. Thus, overall, it appeared that, in zebrafish, temperature is the environmental parameter that most significantly impacted gene expression levels. Notably, both well-defined CNSS genes such as *uts1, uts2a* and *uts2b* and other genes were affected.

**Fig 4.**
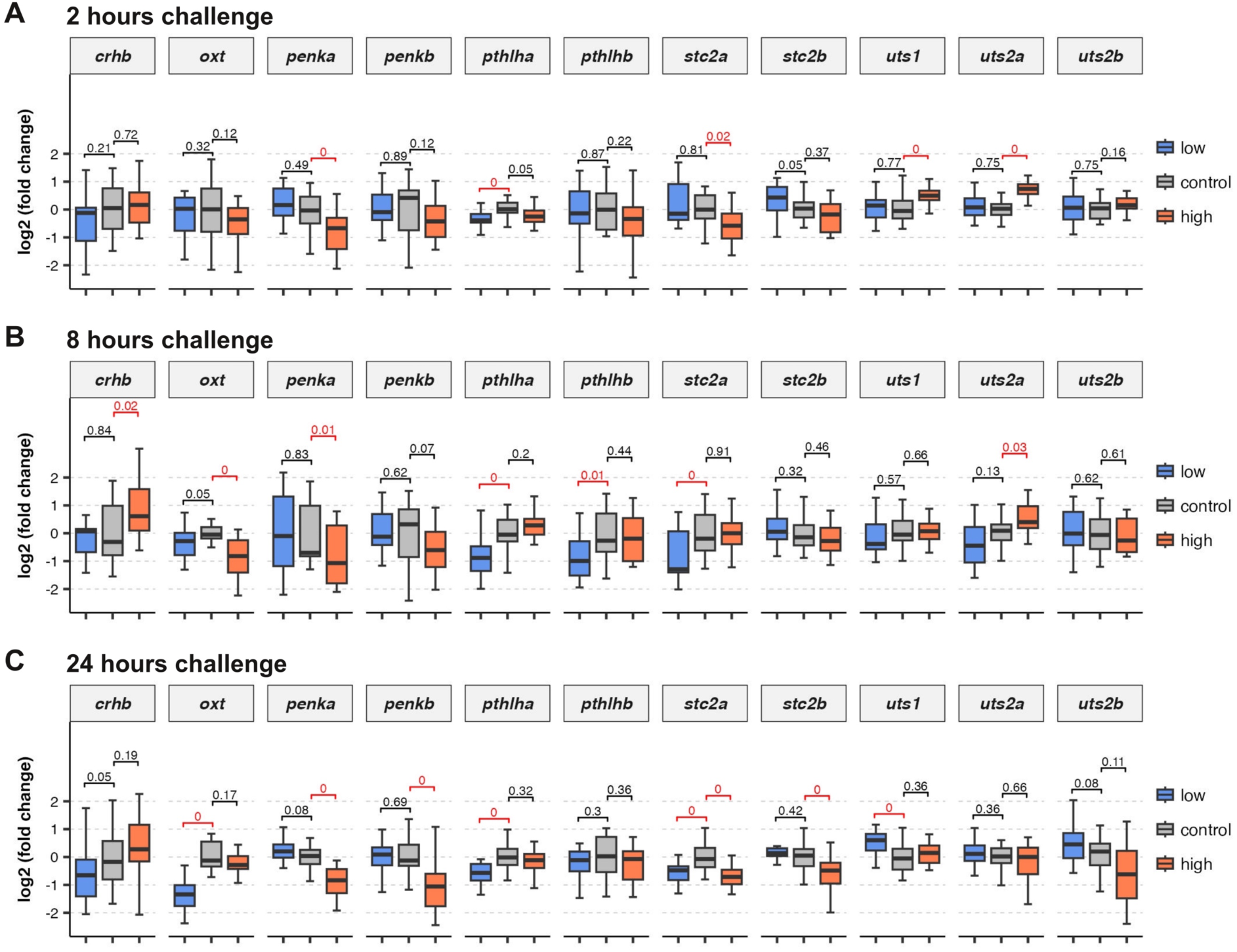
Gene expression changes following acute temperature challenge. Expression levels of indicated genes after 2 hours (A), 8 hours (B) and 24 hours (C) temperature challenge with low, control and high conditions. In all cases, results are expressed as log2 of fold change (ΔΔCT) compared to the control condition.

## 4 Discussion

Although first described 70 years ago (Enami and Imai, 1955), the physiological significance of the CNSS remains quite enigmatic. As a well-established genetic model organism for several decades, the zebrafish could help shed new light on the function of the CNSS. In this study, as a first step in this initiative, we attempted to identify environmental factors whose variations induce changes in CNSS activity. To ensure that the observed effects were physiologically relevant, the variations in water pH, salinity and temperature used in this study, were chosen in the range of conditions that zebrafish can encounter in their natural habitat (Arunachalam et al., 2013; Engeszer et al., 2007). One exception was the high salinity condition, for which we used water with 5 g/l of sea salt, a concentration commonly employed as a treatment for skin infections in freshwater fish. Our results showed that CNSS activity, assayed by expression levels of peptide hormone-encoding genes, is modified upon acute change in water temperature, salinity, and, to a lesser extent, pH. This suggest that, in the zebrafish, the CNSS is involved in osmoregulation and thermo-adaptation, which is consistent with what has been reported in other species (Rousseau et al., 2024), and shows that the zebrafish is a relevant model to study the function of the CNSS.

The implication of CNSS in osmoregulation has been proposed just after its initial description by Enami, who found that osmotic stimuli induced histological changes of the CNSS in the pond loach (*Misgurnus anguillicaudatus*) (Enami, 1956). Since then, several studies have further supported this hypothesis in different species (Rousseau et al., 2024). Yet, in the zebrafish, this had not been clearly shown since the only two reported studies (Jin et al., 2017; Luo et al., 2014) with this model used embryos and young larvae (between 2 and 6 dpf) in which the CNSS is likely not yet fully functional (Fridberg and Bern, 1968). Moreover, in these studies, mRNAs were obtained from entire animals, raising questions about the source of the transcripts. Thus, our work provides new evidence for a role of the CNSS in osmoregulation in zebrafish. Our results also suggested that the CNSS activity is influenced by the pH. In our pH challenge, the overall salinity was identical in all three conditions and only the ionic composition was modified. Although the observed changes in CNSS gene expression were modest, our results suggest that the CNSS is also involved in the physiological adaptation to subtle changes in water composition. This is in line with a study on brook trout (*Salvelinus fontinalis)* that reported that acidified water appears to stimulate release of Uts2 from the urophysis (Chevalier et al., 1986).

Recent studies have suggested that the CNSS is also involved in thermo-adaptation in the olive flounder (*Paralichthys olivaceus*) (Yuan et al., 2021, 2020, 2019). Earlier works in zebrafish had reported that temperature challenges modified expression levels of CNSS genes, but, again, this was done in embryos and young larvae (Jin et al., 2017; Luo et al., 2014). In our study, the temperature challenges resulted in the highest number of genes with modified expression levels. This suggests that, in the zebrafish, the CNSS could be more important for thermo-adaptation than for osmoregulation. Yet, while *uts1, uts2a* and *uts2b* are specifically expressed in Dahlgren cells in the caudal part of the fish (Parmentier et al., 2008), other genes could be expressed in other tissues. Moreover, changes in temperature modify oxygen levels, which could also influence the activity of the CNSS. Thus, further analysis will be needed to confirm the implication of the CNSS in thermo-adaptation in zebrafish.

Because we aimed to assay the effect of several parameters and conditions, we used juvenile zebrafish (10 weeks), which are easier to produce in large numbers compared to fully grown adults. However, the CNSS seems to form relatively late during fish development (Fridberg and Bern, 1968; Oka et al., 1993; Cioni et al., 2000) and, as a consequence, we cannot rule out that in juvenile zebrafish, the CNSS is not yet fully mature. This could result in lower gene expression levels and responses to challenges. Also, because we used animals before sexual maturity, we did not attempt to differentiate males and females. Nevertheless, it is possible that fine differences in expression levels and responses are already present at this stage. This may have increased variability between samples and diminished the power of our assay. It is thus clear that our results will have to be confirmed in adults.

In conclusion, our results provide the first evidence that the CNSS in zebrafish is involved in thermo-adaptation and osmoregulation. The zebrafish stands out as a model organism thanks to its unique potential for functional studies, notably thanks to efficient methods for CRISPR/Cas9-mediated genome editing and for transgenesis. Mutant lines, could be used to study the consequences of genes encoding peptide hormones in control or modified water conditions. A limitation is that some genes studied in this work are also expressed outside of the CNSS, notably in the brain. Identifying Dahlgren cell-specific regulatory sequences of these genes would enable tissue specific gene inactivation (Kalvaitytė and Balciunas, 2022) to precisely investigate their function in the CNSS. Further, mutant of genes encoding for the receptors for these hormones could help to understand the effect of CNSS product on target tissues. Additionally, these regulatory sequences could be harnessed to perform genetic ablation of Dahlgren cells in order to revisit the effect of CNSS ablations (Labbaf et al., 2022; Mathias et al., 2014). Thus, our work contributes to position the zebrafish as model of choice to study the role of the CNSS in fish homeostasis and should pave the way for further studies aimed at understanding the role of this neuroendocrine system, which is unique to fishes.

## Acknowledgments

The authors are grateful to Céline Maurice, Philippe Durand and Jean-Paul Chaumeil (PhyMA, MNHN) for zebrafish care. We thank Fabrice Girardot and Karine Rousseau for helpful discussions and critical reading of the manuscript.

## Author contributions

Conceptualization and methodology: Guillaume Pézeron, Bérénice Bichon and Hervé Tostivint.

Investigation: Bérénice Bichon, with support from Gladys Alfama and Anne-Laure Gaillard.

Formal analysis and validation: Guillaume Pézeron and Bérénice Bichon

Visualization and writing: Guillaume Pézeron and Bérénice Bichon

