## Supplementary material for "Hormone gene expression changes in the zebrafish caudal neurosecretory system under acute environmental challenges": Supplmental Table 1

| **Oligo name** | **Sequence** |
| --- | --- |
| *avp_F* | TCTGCCTGCTACATCCAGAAC |
| *avp_R* | AGATACTGGGGCCAAAACAGC |
| *crha_F* | GGAGGACACCGACTGAGATTAC |
| *crha_R* | AGAGCTCCAGACGGAGAGTC |
| *crhb_F* | CGCGCAAAGTTCAAAAACCATC |
| *crhb_R* | CCGTGGTGACGAGAAAATTGAG |
| *lsm12b_F* | GAGACTCCTCCTCCTCTAGCAT |
| *lsm12b_R* | GATTGCATAGGCTTGGGACAAC |
| *mob4_F* | AAGAGTGCCCTGCCATTGATTA |
| *mob4_R* | AGTTTGGCCACAGATGATTCCT |
| *oxt_F* | CCTTCTCCGTGTGAGATGTCTG |
| *oxt_R* | ATCTCCGTCCACACACGACT |
| *penka_F* | CTCCGACAGACAGACATCGAC |
| *penka_R* | CCTCTTCGGTGAGGATGTTTTTAC |
| *penkb_F* | ATGCCCCTTCAGCTCAAACA |
| *penkb_R* | AATGTTTGCACTGACTCCAAGAG |
| *pthlha_F* | TGCTCTGTTTTTACTGTGTTCTCC |
| *pthlha_R* | CGGCCTTTATCATGCATCAGC |
| *pthlhb_F* | CAATCCCGATACCCATCCAGTC |
| *pthlhb_R* | CTCTTCCGGTCATGCAGAGAG |
| *stc2a_F* | AACGTCAATGTCATGGTGGAGA |
| *stc2a_R* | CGAGTAATGGCTTCCTTCACCT |
| *stc2b_F* | CATGCACTTGAGCGTGTCTTTA |
| *stc2b_R* | GAGATCCGCGGACTACATGAT |
| *tdr7_F* | GCAGCATAATTGAGTACACCC |
| *tdr7_R* | TTGCCTATATTCACTGAGAAATGGA |
| *uts1_F* | AACAGAAGACGATCGAGAAGTGC |
| *uts1_R* | ATGTGGCTCAGTAAGACTGAAGTG |
| *uts2a_F* | TCCTTTCTGCTCATCCCGTTAC |
| *uts2a_R* | ACATCAGTGAGGTCAGATACAGAG |
| *uts2b_F* | GTTACCCGTCTCTCATCAGTGG |
| *uts2b_R* | GTTTTTCCAGCAAGGCCTCTTT |
